# Circulating concentrations of micro-nutrients and risk of breast cancer: A Mendelian randomization study

**DOI:** 10.1101/668186

**Authors:** N. Papadimitriou, N. Dimou, D. Gill, I. Tzoulaki, N. Murphy, E. Riboli, S. J. Lewis, R. M. Martin, M. J. Gunter, K. K. Tsilidis

**Author notes:** Corresponding author: Dr Konstantinos K Tsilidis, Department of Epidemiology and Biostatistics, Imperial College London, St Mary’s Campus, London, W2 1PG, United Kingdom.

## Abstract

**Background:** The epidemiological literature reports inconsistent associations between consumption or circulating concentrations of micro-nutrients and breast cancer risk. We investigated associations between genetically determined concentrations of 11 micro-nutrients (beta-carotene, calcium, copper, folate, iron, magnesium, phosphorus, selenium, vitamin B6, vitamin B12 and zinc) and breast cancer risk using Mendelian randomization (MR).

**Materials and methods:** A two-sample MR study was conducted using 122,977 women with breast cancer, of whom 69,501 were estrogen receptor positive (ER^+ve^) and 21,468 were ER^−ve^, and 105,974 controls from the Breast Cancer Association Consortium. MR analyses were conducted using the inverse variance weighted approach, and sensitivity analyses were conducted to assess the impact of potential violations of MR assumptions.

**Results:** One standard deviation (SD: 0.08 mmol/L) higher genetically determined concentration of magnesium was associated with a 17% (odds ratio [OR]: 1.17, 95% confidence interval [CI]: 1.10 to 1.25, *P*=9.1 × 10^−7^) and 20% (OR: 1.20, 95% CI: 1.11 to 1.30, *P*=3.17 × 10^−6^) higher risk of overall and ER^+ve^ breast cancer, respectively. An inverse association was observed for a SD (0.5 mg/dL) higher genetically determined phosphorus concentration and ER^−ve^ breast cancer (OR: 0.84, 95% CI: 0.72 to 0.98, *P*=0.03). A suggestive inverse association was observed for a SD (0.48 mg/dL) higher genetically determined calcium concentration with overall breast cancer (OR: 0.91, 95% CI: 0.83 to 1.00, *P*=0.06). There was little evidence that any of the other nutrients were associated with breast cancer. The results for magnesium were robust under all sensitivity analyses.

**Conclusions:** Higher circulating concentrations of magnesium, phosphorus and calcium may affect breast cancer risk. Further work is required to replicate these findings and investigate underlying mechanisms.

**key message:** We conducted a Mendelian randomization study to investigate whether concentrations of 11 micro-nutrients are associated with risk of breast cancer. An increased risk of overall and oestrogen-receptor positive disease was observed for genetically higher concentrations of magnesium and inverse associations were observed for phosphorus and calcium concentrations

Where authors are identified as personnel of the International Agency for Research on Cancer / World Health Organization, the authors alone are responsible for the views expressed in this article and they do not necessarily represent the decisions, policy or views of the International Agency for Research on Cancer / World Health Organization.

## Introduction

Breast cancer is the most common cancer in women with an estimated 2.1 million new cases and over half a million deaths worldwide in 2018 [1]. Breast cancer is a heterogeneous disease and its epidemiology varies with menopausal and tumour receptor status at time of diagnosis. Established risk factors for post-menopausal breast cancer include adiposity, endogenous concentrations of estrogens and androgens and reproductive factors such as early menarche, late menopause, late age at first pregnancy and hormone replacement therapy [2]. The role of nutrition in breast cancer development is unclear. The World Cancer Research Fund (WCRF) has recently systematically reviewed all literature on the associations of dietary and nutritional factors with breast cancer [3]. The WCRF report concluded that the evidence was strong only for a positive association between alcohol consumption and risk of pre- and post-menopausal breast cancer, whereas limited but suggestive evidence was found for inverse associations between consumption of non-starchy vegetables and foods containing carotenoids and calcium with risk of pre- and post-menopausal breast cancer [3]. The literature on circulating concentrations of minerals and vitamins with risk of breast cancer is generally not extensive with perhaps the exceptions of carotenoids and vitamin D [3].

Most of the evidence regarding the nutritional epidemiology of breast cancer comes from observational studies that often rely on food frequency questionnaires (FFQ) to measure the consumption of foods and nutrients. This approach is prone to measurement error, because it is based on participants’ self-reports that might be inaccurate leading to biased results [4]. Furthermore, individuals who follow different dietary patterns might also differ in other aspects, which are not always adequately controlled for in statistical confounder adjustments [5, 6]. Evidence from clinical trials is also lacking, as there are not many adequately powered trials that have evaluated the effect of micro-nutrients and cancer risk [7-10].

A Mendelian randomization (MR) approach can be used to assess the existence of a potential causal association between a risk factor and disease. MR attempts to emulate a randomized controlled trial (RCT) within an observational setting using genetic variants as instrumental variables for the risk factor of interest [11]. As genotype is allocated randomly at conception, genetic variants are not influenced by potential confounding factors, such as environmental exposures, and cannot be altered by disease occurrence.

The aim of the current study was to investigate potential causal associations of genetically determined circulating concentrations of minerals and vitamins with risk of overall breast cancer and cancer by estrogen receptor status (ER^+ve^ and ER^−ve^) using MR methodology.

## Methods

### Data on the genetic epidemiology of circulating nutrients concentrations

The Genome-Wide Association Studies (GWAS) catalog (https://www.ebi.ac.uk/gwas) and Pubmed (https://www.ncbi.nlm.nih.gov/pubmed) were searched (last checked on September 2017) for published GWASs on circulating concentrations of minerals and vitamins that were conducted in populations of European ancestry. The initial list included 20 nutrients: beta-carotene, calcium, copper, folate, iron, magnesium, phosphorus, potassium, retinol, selenium, sodium, vitamins B1, B2, B6, B12, C, D, E, and K, and zinc. Vitamin D was excluded because recent MR studies have investigated the role of circulating vitamin D concentrations and risk of breast cancer [12, 13]. Potassium, sodium, vitamins B1, B2, C and K were also excluded because either no genome-wide significant results have been reported [14, 15] or no GWAS has been conducted. Two further GWAS, those for circulating vitamin E and retinol, were excluded because they adjusted for body mass index (BMI), which may bias MR estimates [16-19]. Published GWAS for 11 nutrients were finally retrieved, namely beta-carotene, calcium, copper, folate, iron, magnesium, phosphorus, selenium, vitamins B6 and B12 and, zinc [14, 20-27]. Single nucleotide polymorphisms (SNPs) that independently affected the concentrations of these nutrients at a genome wide significance level (*p* < 5 × 10^−8^) and were not in linkage disequilibrium (LD r^2^ ≤ 0.1) were obtained. For serum iron, we used summary estimates for three (rs1800562, rs1799945 and rs855791) out of the five (rs1800562, rs1799945 and rs855791, rs7385804, rs8177240) available genome-wide significant SNPs, because these SNPs showed a concordant effect on serum iron, ferritin, transferrin and transferrin saturation, and have been associated with an overall increased systemic iron status [26, 28, 29]. Three additional SNPs with minor allele frequency (MAF) smaller than 5% (rs12272669, rs2336573, rs6859667) in the initial GWAS papers for selenium and vitamin B12 were excluded from the final list of SNPs since their effects might be unstable due to their small MAF [24, 25]. In total, summary association data for 47 common (minor allele frequency ≥ 0.05) genetic variants associated with the 11 nutrients of interest was obtained. Detailed information on the selected genetic variants is provided on Supplementary Table S1.

### Data on the genetic epidemiology of breast cancer

The genetic effects of the selected instruments on risk of overall breast cancer and cancer by estrogen receptor status were obtained from a recently published large GWAS of almost 230,000 participants (122,977 breast cancer cases and 105,974 controls) of European ancestry from the Breast Cancer Association Consortium (BCAC) (http://bcac.ccge.medschl.cam.ac.uk/) [30]. Out of the 47 SNPs, rs778805, that was associated with vitamin B12, was significantly associated with risk of overall breast cancer, and rs4072037, that was associated with magnesium, was significantly associated with both overall and ER^+ve^ breast cancer (Supplementary Table S1).

### Statistical power

Power calculations were performed using an online tool available at http://cnsgenomics.com/shiny/mRnd/ [31]. The statistical power to capture an OR of 1.10 per a standard deviation (SD) change in the circulating concentrations of the nutrients, given a sample size of 228,951 participants, type 1 error of 5% and a proportion of cases equal to 0.54, ranged from 0.55 for calcium to 1 for copper, vitamin B12, and, zinc and the statistical power was between 0.80 and 1 for seven (i.e., beta – carotene, copper, iron, magnesium, selenium, vitamin B12, and zinc) of the 11 instruments tested (Supplementary Table S2).

### Mendelian Randomization analysis

Two-sample MR using summary association data was performed. In the case of beta-carotene, where only one SNP was available, the effect estimate was calculated as the ratio of the SNP-outcome over the SNP-nutrient association [32], whereas the fixed – effect inverse variance weighted (IVW) method was implemented when the instruments consisted of multiple SNPs, and this analysis can be thought as a meta-analysis of single SNP effects [32, 33]. The beta estimates from the regressions for circulating concentrations for beta-carotene, copper, selenium, vitamin B6, and zinc were transformed from the logarithmic scale that were originally reported in the published GWAS to the natural scale using a published formula [34]. All reported associations correspond to OR for risk of breast cancer per SD change in the circulating concentrations of the nutrients.

To be valid instruments for the MR analysis, the selected genetic variants must meet the following criteria: i) be associated with the circulating concentrations of the nutrients, ii) be independent of any potential confounding variable of the nutrient – breast cancer association, and iii) affect breast cancer only through the nutrient being instrumented and not via any other pathway [35]. The strength of each instrument was measured using the F statistic calculated by the formula: *F* = *R*^2^(*N* − 2)/(1 − *R*^2^), where R^2^ is the proportion of the explained variance of the nutrient by the genetic instrument and *N* the sample size of the GWAS for the SNP-nutrient association [36]. The R^2^ was calculated using an already published formula [37].

### Sensitivity analyses

The second and third MR assumptions was examined by performing the following statistical analyses: Cochran’s Q [38], random effects IVW MR analysis [39], MR-Egger regression [40], weighted median [35], weighted mode approach (WMA) [41], the MR pleiotropy residual sum and outlier test (MR-PRESSO) [42], and a leave one SNP out analysis. More details are provided in the supplementary methods. We further evaluated whether the selected genetic instruments were associated with secondary phenotypes in Phenoscanner (http://www.phenoscanner.medschl.cam.ac.uk/) and GWAS catalog (last checked May 2018) [43, 44]. All analyses were implemented in the statistical software R version 3.4.3 using the package MendelianRandomization [45].

## Results

### Mendelian randomization estimates

A SD (0.08 mmol/L) higher genetically determined concentration of magnesium was associated with a 17% (OR: 1.17, 95% CI: 1.10 to 1.22, *P*=9.10×10^-7^) and 20% (OR: 1.20, 95% CI: 1.11 to 1.30, *P*=3.17×10^-6^) higher risk of overall and ER^+ve^ breast cancer, respectively. An inverse association was observed for a SD (0.50 mg/dL) higher genetically determined phosphorus concentration and risk of ER^−ve^ breast cancer (OR: 0.84, 95% CI: 0.72 to 0.98, *P*=0.03). A suggestive inverse association was observed for a SD (0.48 mg/dL) higher genetically determined calcium concentration and risk of overall breast cancer (OR: 0.91, 95% CI: 0.83 to 1.00, *P*=0.06). There was little evidence that any of the other nutrients were associated with risk of breast cancer or its subtypes (Figure 1).

**Figure 1.**
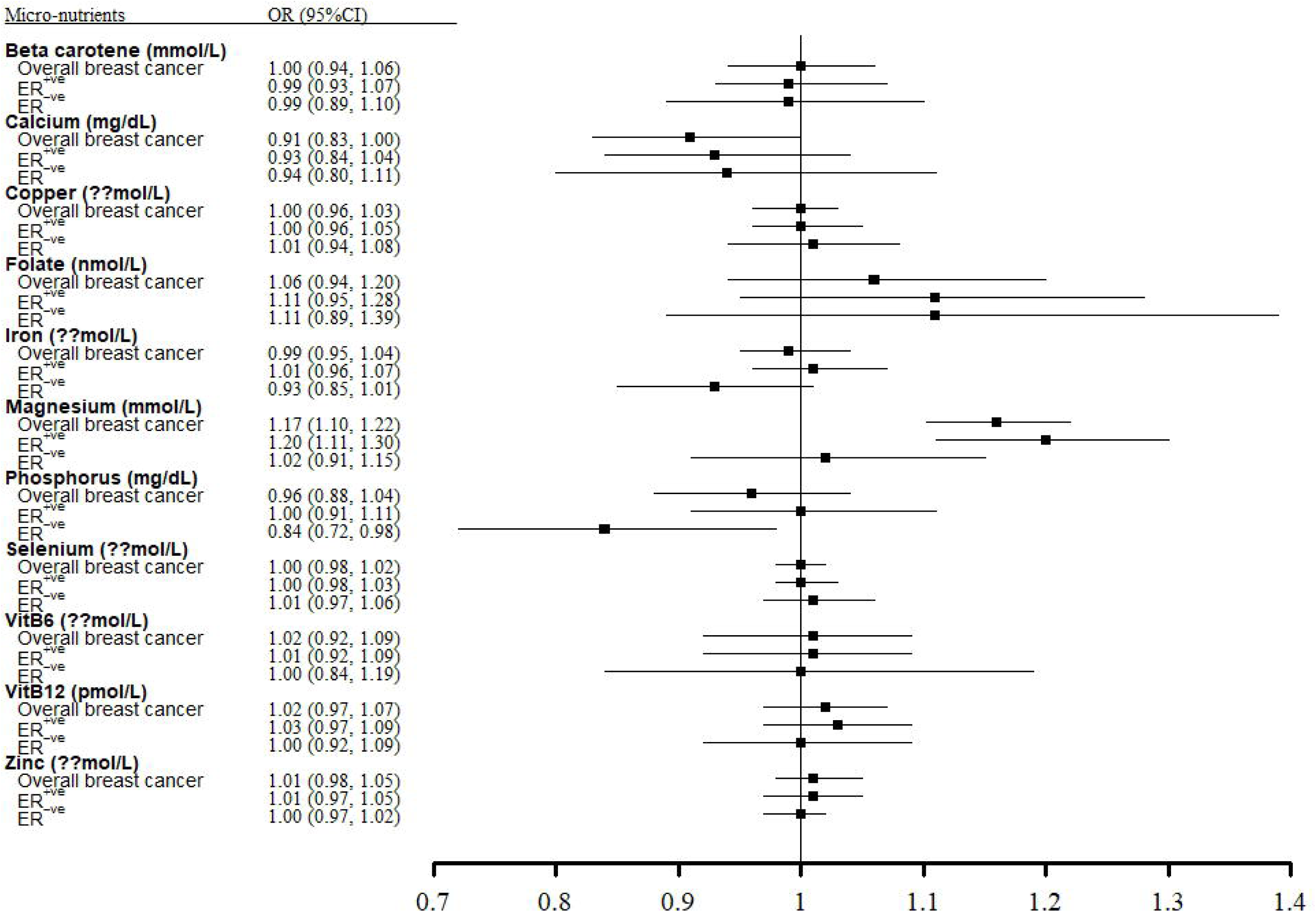
Forest plot showing results from the Mendelian randomization study to evaluate potential causal associations between 11 nutrients and breast cancer both overall and by estrogen receptor status. The odds ratios (OR) were calculated using the inverse variance weighted (IVW) method and correspond to a 1-SD increase in the concentration of the nutrients.

### Evaluation of Mendelian randomization assumptions

The F-statistic was larger than 10 for all the included variants implying absence of weak instruments (minimum value=16, maximum value=420) (Supplementary Table S1).

Cochran’s Q test was not significant for all analyses, except for vitamin B12 and overall breast cancer risk (Cochran’s Q *P*= 0.02) however, no change was observed in the results after performing the random effects IVW MR (Supplementary Table 3). Additionally, some indication of horizontal pleiotropy was found in the analysis of calcium and overall breast cancer risk based on the MR-Egger test (intercept *P*= 0.02) (Supplementary Table S3).

The MR-Egger regression, the weighted median and the weighted mode methods confirmed the IVW per a SD higher genetically determined magnesium concentrations for overall breast cancer (*OR*_*MR*−*Egger*_=1.24, 95% CI: 1.01 to 1.53; *OR*_*weighted median*_=1.20, 95% CI: 1.10 to 1.31, *OR*_*MBE*_ =1.21, 95% CI: 1.12 to 1.39) and ER^+ve^ breast cancer (OR_*MR*−*Egger*_ =1.44, 95% CI: 1.05 to 1.97; *OR*_*weighted median*_ =1.19, 95% CI: 1.05 to 1.34, *OR*_*weighted mode*_ =1.36, 95% CI: 1.16 to 1.59) (Supplementary Table S4). A positive association was observed between genetically determined vitamin B12 concentrations and overall breast cancer risk based on the weighted median approach (OR: 1.08, 95% CI: 1.01 to 1.15). There was little evidence that any of the other nutrients were associated with risk of breast cancer in sensitivity analyses (Supplementary Table S4). Finally, the MR-PRESSO analysis did not reveal any outlying SNPs, which was also evident when we estimated and plotted the MR results for all nutrients by each separate SNP (Supplementary Figures S1-S10).

When we iteratively excluded one SNP at a time from all genetic instruments to further probe into potential SNP outliers, the positive association between genetically determined magnesium concentrations and risk of overall and ER^+ve^ breast cancer remained significant (Supplementary Table S5). However, when rs4072037 was removed from the genetic instrument for magnesium concentrations, the associations for risk of overall (OR: 1.11, 95% CL: 1.02 to 1.21) and ER^+ve^ breast cancer (OR: 1.09, 95% CL: 0.98 to 1.21) were slightly attenuated. There was little evidence that any of the other nutrients were associated with risk of breast cancer in the leave one out analysis with the exceptions described below. The inverse association observed in the IVW analysis between genetically determined phosphorus concentration and the risk of ER^−ve^ breast cancer was not present after removal of rs1697421, rs947583 or rs297818. When rs1801725 was excluded from the genetic instrument of calcium concentrations, a protective effect was observed for overall (OR: 0.79, 95% CI: 0.68 to 0.91) and ER^+ve^ breast cancer (OR: 0.82, 95% CL: 0.69 to 0.98).

Several genome-wide significant associations of the genetic instruments with secondary traits were observed (Supplementary Table S6). Some of these may be associated with breast cancer and are potential causes of horizontal pleiotropy, namely associations with diabetes, height, insulin resistance, C-reactive protein, glycosylated haemoglobin, and age at menarche. However, when the SNPs that were associated with the latter secondary phenotypes were excluded in the leave one out analysis, the observed associations of the genetic instruments with breast cancer did not change (Supplementary Table S5).

## Discussion

In this comprehensive MR analysis of 11 nutrient concentrations with risk of breast cancer, a positive association was observed for genetically elevated concentrations of circulating magnesium and risk of total and ER^+ve^ breast cancer, which was robust to sensitivity analyses. Potential inverse associations of circulating phosphorus and calcium concentrations with ER^−ve^ and overall breast cancer, respectively, were observed in the main analysis, but were not robust to sensitivity analyses.

The literature on the association of dietary or circulating magnesium and the risk of breast cancer is scarce and inconclusive. Only one cohort study, involving 12,902 women aged 35-64 years at baseline with 415 breast cancers developed during a mean follow-up of 11 years, has reported on dietary magnesium in association with overall breast cancer risk. This study reported little evidence for an association (hazard ratio [HR]: 0.97, 95% CI: 0.69 to 1.35) comparing the 1^st^ to the 4^th^ quartile of magnesium consumption [46]. In a cross-sectional study of 725 women, urinary magnesium levels were positively associated with breast density, although the interpretation of this finding is unclear, as breast density is a risk factor for breast cancer but it may also obscure detection of the disease [47-49]. The genetic instruments used in our analysis affect magnesium homeostasis and mechanistic studies suggest that increased concentrations of magnesium within breast cancer cells can lead to tumour progression through the regulation of enzymes involved in energy generation, and its presence is also needed for cell adhesion and cancer metastasis [50, 51]. Evidence from animal studies suggests that magnesium has a protective effect in the early phases of chemical carcinogenesis but promotes tumour growth [52]. There are no magnesium supplementation trials assessing relevant cancer outcomes. Our finding of a positive association between genetically determined magnesium concentrations and risk of breast cancer should be further evaluated in humans and animals using observational and interventional designs.

An inverse association was observed in our analysis between concentrations of phosphorus and ER^−ve^ breast cancer. We found only one relevant cohort study involving 186,620 women, where 4,925 incident breast cancers developed during a mean follow-up of 13 years, reporting that a SD (0.17 mmol/L) increase in serum inorganic phosphate levels were inversely associated with total breast cancer risk (HR: 0.93, 95% CI: 0.90 to 0.96), but these estimates were not adjusted for potentially important confounders such as alcohol consumption and smoking status [53]. Our analysis did not confirm this association for total breast cancer (OR: 0.96, 95% CI: 0.88 to 1.04), but found an inverse association for ER^−ve^ disease (OR: 0.84, 95% CI: 0.72 to 0.98). Studies have linked estrogen levels with negative regulation of circulating inorganic phosphate and high levels of estrogen are strongly positively associated with breast cancer [54], but this mechanism is more relevant for ER^+ve^ than ER^−ve^ breast cancer.

Our primary analysis suggested an inverse association of calcium with breast cancer (OR: 0.91, 95% CI: 0.83 to 1.00). After the exclusion of rs1801725, this association stregthened for overall (OR: 0.79, 95% CI: 0.68 to 0.91) and ER^+ve^ breast cancer (OR: 0.82, 95% CI: 0.69 to 0.98). Rs1801725 is located in the calcium-sensing receptor (CaSR) gene, which plays an important role in calcium metabolism and has been suggested to act as an oncogene for several cancers including breast [55]. However, results from GWAS indicate that rs1801725 is also associated with immunoglobulin E (IgE), bone mineral density, childhood obesity and adiponectin concentrations (all *P*<0.005) [56-59], which might reflect pleiotropic actions of this genetic variant. A recent meta-analysis of three prospective studies also reported an inverse association between serum calcium and breast cancer risk (RR: 0.80, 95% CI: 0.66 to 0.97, per 1 mmol/L increase in serum calcium) [60]. In addition, the third expert report from WCRF also reported an inverse association of dietary calcium with pre-menopausal (relative risk [RR] for 300mg/day increase of dietary calcium intake: 0.87, 95% CI: 0.76 to 0.99) and post-menopausal breast cancer (RR: 0.96, 95% CI: 0.94 to 0.99), but the evidence has been graded as limited-suggestive [3]. Possible mechanistic explanations could be that increased cytosolic levels of calcium might affect cell cycle processes and apoptosis through Ras and β-catenin pathways [61, 62].

Our main analysis showed little evidence of an association between genetically determined circulating vitamin B12 concentrations with risk of breast cancer, although one of the vitamin B12 SNPs, rs778805, was associated with risk of overall breast cancer in the BCAC GWAS and the result of the weighted median analysis was significant for overall breast cancer. A recent meta-analysis of four observational studies concluded that there was no association between serum vitamin B12 and the risk of overall breast cancer (RR: 0.73, 95% CI: 0.44 to 1.22) comparing the highest vs lowest categories [63].

The main strengths of the current MR study are the ability to circumvent biases that often plague observational studies and the utilization of data from large-scale genetic consortia. Potential limitations should be also considered in the interpretation of our findings. Conducting an MR analysis using summary association data does not allow for stratified analyses by covariates of interest such as menopausal status and body mass index. Moreover, our analysis was underpowered to identify associations of small magnitude between some nutrients and risk of breast cancer. Specifically, a minimum R^2^ value of 1.5% was required in our study to capture an OR of 1.10 per SD increase in nutrient concentrations with 0.8 power, and this cutoff point was not met for folate (R^2^ =1%), vitamin B6 (1%), calcium (0.8%) and phosphorus (1.2%). Larger GWASs of nutrients are required to enable the construction of better genetic instruments for these nutrients. Some of the sensitivity analyses such as the MR-Egger regression are underpowered when a small number of genetic instruments are used. We did not correct for multiple comparisons, but if a more conservative significance threshold based on Bonferroni correction (0.05/11=0.0045) would have been applied, then only the results for magnesium would remain statistically significant.

In conclusion, we conducted the first comprehensive two-sample MR study to investigate whether genetically determined concentrations of 11 micro-nutrients are associated with risk of breast cancer. An increased risk of overall and ER^+ve^ disease was observed for genetically higher concentrations of magnesium that was robust to several different sensitivity analyses. Future studies are warranted to replicate this finding and to disentangle the potential underlying pathways.

## Acknowledgements

The breast cancer genome-wide association analyses were supported by the Government of Canada through Genome Canada and the Canadian Institutes of Health Research, the ‘Ministère de l’Économie, de la Science et de l’Innovation du Québec’ through Genome Québec and grant PSR-SIIRI-701, The National Institutes of Health (U19 CA148065, X01HG007492), Cancer Research UK (C1287/A10118, C1287/A16563, C1287/A10710) and The European Union (HEALTH-F2-2009-223175 and H2020 633784 and 634935). All studies and funders are listed in Michailidou et al (2017).

We would also like to thank Professor David Evans and Professor John Whitfield for allowing us the access to the complete list of results for selenium from their GWAS study, as well as, with the additional information on the mean and standard deviation values of the three minerals (copper, selenium, zinc) required to transform the results from the logarithmic to the natural scale.

---

**Supplementary Figure S1: Forest plots showing MR analyses by each SNP separately and the pooled MR result for calcium and breast cancer risk, overall and by estrogen receptor status**. The odds ratios (OR) were calculated using the inverse variance weighted (IVW) method and correspond to a 1-SD increase in the concentration of calcium.

**Supplementary Figure S2: Forest plots showing MR analyses by each SNP separately and the pooled MR result for copper and breast cancer risk, overall and by estrogen receptor status**. The odds ratios (OR) were calculated using the inverse variance weighted (IVW) method and correspond to a 1-SD increase in the concentration of copper.

**Supplementary Figure S3: Forest plots showing MR analyses by each SNP separately and the pooled MR result for folate and breast cancer risk, overall and by estrogen receptor status**. The odds ratios (OR) were calculated using the inverse variance weighted (IVW) method and correspond to a 1-SD increase in the concentration of folate.

**Supplementary Figure S4: Forest plots showing MR analyses by each SNP separately and the pooled MR result for iron and breast cancer risk, overall and by estrogen receptor status**. The odds ratios (OR) were calculated using the inverse variance weighted (IVW) method and correspond to a 1-SD increase in the concentration of iron.

**Supplementary Figure S5: Forest plots showing MR analyses by each SNP separately and the pooled MR result for magnesium and breast cancer risk, overall and by estrogen receptor status**. The odds ratios (OR) were calculated using the inverse variance weighted (IVW) method and correspond to a 1-SD increase in the concentration of magnesium.

**Supplementary Figure S6: Forest plots showing MR analyses by each SNP separately and the pooled MR result for phosphorus and breast cancer risk, overall and by estrogen receptor status**. The odds ratios (OR) were calculated using the inverse variance weighted (IVW) method and correspond to a 1-SD increase in the concentration of phosphorus.

**Supplementary Figure S7: Forest plots showing MR analyses by each SNP separately and the pooled MR result for selenium and breast cancer risk, overall and by estrogen receptor status**. The odds ratios (OR) were calculated using the inverse variance weighted (IVW) method and correspond to a 1-SD increase in the concentration of selenium.

**Supplementary Figure S8: Forest plots showing MR analyses by each SNP separately and the pooled MR result for vitamin B6 and breast cancer risk, overall and by estrogen receptor status**. The odds ratios (OR) were calculated using the inverse variance weighted (IVW) method and correspond to a 1-SD increase in the concentration of vitamin B6.

**Supplementary Figure S9: Forest plots showing MR analyses by each SNP separately and the pooled MR result for vitamin B12 and breast cancer risk, overall and by estrogen receptor status**. The odds ratios (OR) were calculated using the inverse variance weighted (IVW) method and correspond to a 1-SD increase in the concentration of vitamin B12.

**Supplementary Figure S10: Forest plots showing MR analyses by each SNP separately and the pooled MR result for zinc and breast cancer risk, overall and by estrogen receptor status**. The odds ratios (OR) were calculated using the inverse variance weighted (IVW) method and correspond to a 1-SD increase in the concentration of zinc.

